# Subcellular Redox Responses Reveal Different Cu-dependent Antioxidant Defenses between Mitochondria and Cytosol

**DOI:** 10.1101/2022.07.21.500983

**Authors:** Yuteng Zhang, Meng-Hsuan Wen, Guoting Qin, Chengzhi Cai, Tai-Yen Chen

**Author notes:** Corresponding author: Tai-Yen Chen.

## Abstract

Excess intracellular Cu perturbs cellular redox balance and thus causes diseases. However, the relationship between cellular redox status and Cu homeostasis and how such an interplay is coordinated within cellular compartments has not yet been well established. Using combined approaches of organelle-specific redox sensor Grx1-roGFP2 and non-targeted proteomics, we investigate the real-time Cu-dependent antioxidant defenses of mitochondria and cytosol in live HEK293 cells. The Cu-dependent real-time imaging experiments show that CuCl_2_ treatment results in increased oxidative stress in both cytosol and mitochondria. In contrast, subsequent Cu depletion by BCS, a Cu chelating reagent, lowers oxidative stress in mitochondria but causes even higher oxidative stress in the cytosol. The proteomic data reveal that several mitochondrial proteins, but not cytosolic ones, undergo significant abundance change under Cu treatments. The proteomic analysis also shows that proteins with significant changes are related to mitochondrial oxidative phosphorylation and glutathione synthesis. The differences in redox behaviors and protein profiles in different cellular compartments reveal distinct mitochondrial and cytosolic response mechanisms upon Cu-induced oxidative stress. These findings provide insights into how redox and Cu homeostasis interplay by modulating specific protein expressions at the subcellular levels, shedding light on understanding the effects of Cu-induced redox misregulation on the diseases.

**Graphical abstract:** 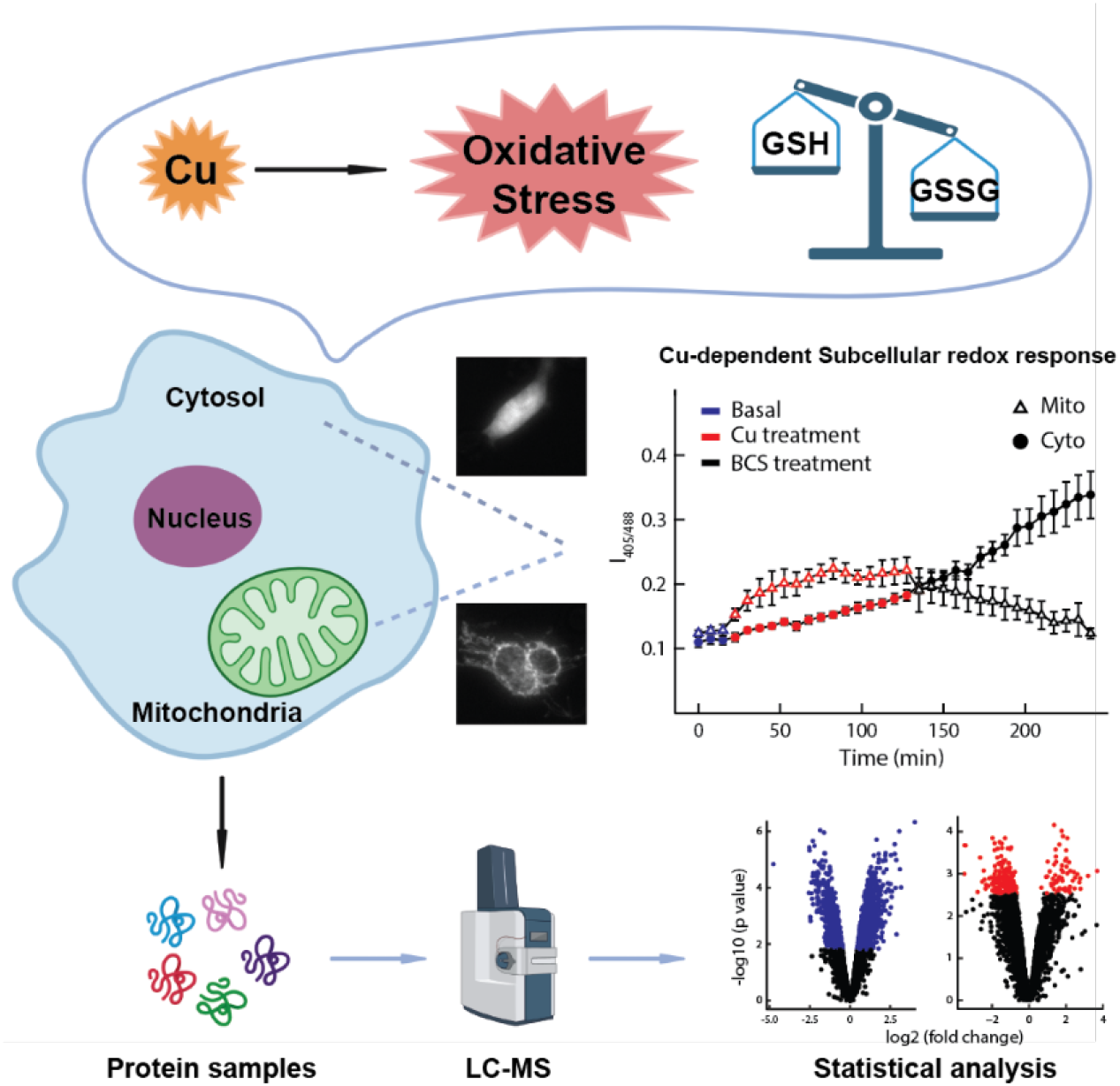

## Introduction

Copper (Cu), as a catalytic and structural cofactor, constitutes active sites of many metalloproteins to enable electron transfer, removal of reactive oxygen species, production of neurotransmitters [1, 2], and cell differentiation [3, 4]. Although essential, Cu’s redox properties also make it detrimental when dysregulated [5, 6]. Misregulation of Cu homeostasis can cause severe damage by producing reactive oxygen species (ROS). Excess Cu ions will react with cellular thiols and oxygen, resulting in the consumption of antioxidant thiols and the generation of reactive oxygen species. Such oxidative stress will obstruct cell proliferation and differentiation, cause apoptosis, damage proteins, lipids, and DNA/RNA, and, therefore, is connected to cancers and neurodegenerative diseases.

The reduced glutathione, GSH, is one of the most important ROS scavengers to provide proper cellular antioxidant defense. Glutathione peroxidase (GPx) and glutaredoxin (Grx) respectively catalytically reduce ROS and oxidatively modified protein thiols at the expense of oxidizing GSH to its oxidized form glutathione disulfide (GSSG), making GSH/GSSG balance a useful measure of oxidative stress [7, 8]. In cells, most GSH resides in the cytosol (∼85%) and mitochondria (∼10%), highlighting that antioxidant defense is specifically crucial in these two compartments [9, 10]. Interestingly, changes in cellular redox status can modulate the redox state of metalloproteins and thus lead to Cu redistribution to different cellular compartments [3, 11]. However, a systematic examination of the effect of Cu on cellular redox status in different compartments remains unavailable. Further, considering their higher Cu levels but lower GSH pool, mitochondria are likely to have different antioxidant defense mechanisms than the cytosol. Identification of the proteins undergoing significant change of abundance in response to Cu induced oxidative stress in the two compartments will shed light on mechanism of subcellular antioxidant defenses.

Several fluorescent probes for sensing biological redox state have been developed and provide advantages, including fast-response, low cost, real-time imaging, and high spatial resolution. For example, Remington’s group reported a ratiometric redox-sensitive green fluorescent protein sensor, roGFP. These GFP-derived biosensors include two cysteines that reversibly switch between sulfanyl group (SH) and disulfide bond (SS) under reduced and oxidized local environments, respectively. Such transition leads to excitation spectral changes and enables redox sensing through fluorescence-ratio microscopy [12]. Dick’s group further fused human glutaredoxin-1 (Grx1) to roGFP2 to generate rapidly reacting glutathione sensor Grx1-roGFP2. Grx1 facilitates specific real-time equilibration between the roGFP2 and the glutathione redox couple (*t*_1/2_ of ∼10 s), allowing dynamic live imaging of the glutathione redox potential in different cellular compartments [7]. On the other hand, liquid chromatography-mass spectrometry (LC-MS) with non-targeted analysis [13, 14] has become the most versatile analytical tool to identify unanticipated metabolites or proteins. The non-targeted analysis allows the detection of hundreds or thousands of molecules simultaneously, enabling the large-scale genome and gene function analysis. In fact, non-targeted multi-omics profiling has successfully identified the novel regulators of mitochondrial cytochrome c oxidase activity [15], mutations promoting cancer cell proliferation [16], the key mechanism for cardiac maturation [17], crosstalk between the SARS-CoV and host [18], and the novel lactones for bacterial quorum-sensing signaling [19].

Here, using combined approaches of organelle-specific redox sensor Grx1-roGFP2 and non-targeted proteomics, we investigate the real-time Cu-dependent antioxidant defenses of mitochondria and cytosol in live HEK293 cells. The time-dependent subcellular redox results revealed that subcellular Cu-induced oxidative stress happens with much slower rates and fewer extents than H_2_O_2_. Mitochondria appear to have more rapid Cu-dependent GSH/GSSG homeostasis than the cytosol. The corresponding proteomics data show that several mitochondrial proteins involved in mitochondrial oxidative phosphorylation and GSH synthesis rapidly undergo significant abundance changes after prolonged Cu treatments. In contrast, cytosolic proteins involved in GSH synthesis undergo substantial changes after Cu reduction. In other words, mitochondria and cytosol have different antioxidant defense mechanisms upon Cu-dependent oxidative stress despite both modulating GSH/GSSG system. Mitochondria respond to Cu treatments and modify protein profile earlier than the cytosol. These findings provide information on how redox status changes within cellular compartments and triggers organelle-specific protein abundance changes in response to Cu-level changes, highlighting the interplay mechanisms between redox and Cu homeostasis.

## Materials and methods

### Plasmids and chemicals

Cytosolic and Mitochondrial redox-sensitive plasmids, mito-Grx1-roGFP2 (#64977) and cyto-Grx1-roGFP2 (#64990) were purchased from Addgene, MA, USA. Hydrogen Peroxide 30% (H_2_O_2_, 1072090250), Dithiothreitol (DTT, D0632-1G), Copper chloride (CuCl_2_, C3279-100G), and bathocuproine sulfonate (BCS, 146625-1G) were purchased from Sigma-Aldrich, St. Louis, MO, USA.

LC-MS grade water, acetonitrile (ACN), formic acid, sequencing grade trypsin, and C18 Ziptips were purchased from Thermo Fisher Scientific (Pittsburgh, PA, USA). All other chemicals were purchased from Millipore Sigma (St. Louis, MO) and used without further purification unless noted otherwise.

### Cell culture and DNA transfection

HEK293 cells (CRL-1573, ATCC) were cultured in Dulbecco’s modified Eagle’s medium (#11965126, Gibco) containing 10% fetal bovine serum (12306C, SAFC), GluataMAX (35050061, Gibco), and Sodium Pyruvate (11360070, Gibco) at 37 °C in 5% CO_2_. Before transfection, cells were seeded into a 24-well plate and grew to a density of 5×10^4^ cells/cm^2^ in each well. The Grx1-roGFP2 plasmid was introduced into cells the next day using Lipofectamine 2000 (Invitrogen) according to the protocol described previously for 36 hours [20]. After transfection, cells were trypsinized and seeded into an imaging chamber for 24-48 hours to reach a cell density of 5×10^4^ cells/cm^2^ before imaging.

### Cell staining

For the colocalization experiment, HEK293 cells were transfected with mito-Grx1-roGFP2 as described above. Mitochondria were counterstained using MITO-ID Red Detection Kit (Cat# ENZ-51007-0100, Enzo Life Sciences) following the manufacturer’s instruction. Briefly, cells were incubated with labeling solution for 15 min at 37 °C in the dark, followed by washing with 100 μL of 1x assay buffer to remove excess reagent. The stained cells were then imaged under a wild-field fluorescence microscope (CKX53, Olympus).

### Real-time redox measurements

All micrographs at different treatment times were collected on an inverted microscope (IX 83, Olympus) using a 40× objective (N1492800, Olympus). The Grx1-roGFP2 transfected samples were excited sequentially with the 405 nm and 488 nm laser (Coherent) under epi-illumination and recorded the GFP emission (emission peak centered around 520 nm) at each timestamp. The final tube lens focused and sent the light through a 525/50 nm bandpass filter (Lot# 344306, Chroma) before forming the final image on the scientific CMOS camera (Prime 95B, Photometrics). Fluorescence intensities of 405 and 488 channels were collected under circularly polarized 405 and 488nm laser with epi excitation and power density around 63 W/cm^2^. Laser power density has been carefully chosen to ensure that the decreases in fluorescence intensities were not due to the photobleaching of the redox sensors.

We have paid close attention to the cell morphology and the transfection level of the redox sensor during the imaging experiments. To ensure our scientific conclusions are physiologically relevant and distilled from robust signals, we only used cells showing clear membrane boundary and nucleus and vigorous Grx1-roGFP2 fluorescent intensity in the bright-field (BF) and fluorescence micrographs, respectively. The mito-Grx1-roGFP2 and cyto-Grx1-roGFP2 were utilized to probe the GSH/GSSG ratio in mitochondria and cytosol, respectively. The fluorescence intensities of all cells were further corrected by subtracting the background reading using a cell-free area in each micrograph for subsequent redox responses analysis under various treatments. 200 μM H_2_O_2_ and 1 mM DTT were used as oxidative and reductive reagents to induce cellular redox responses. For Cu-dependent redox measurements, CuCl_2_ and BCS with a concentration of 200 μM were employed as Cu-stressing and Cu-depletion reagents, respectively. The entire imaging records cellular redox responses under basal, oxidative /Cu-stressed, and reductive/Cu-depleted conditions.

### Quantification of GSH/GSSG balance through ratiometric imaging analysis

To quantify the GSH/GSSG balance in mitochondria and cytosol, we analyzed all fluorescence micrographs using a home-built MatLab (The MathWorks, www.mathworks.de) code. The code is comprised of three major steps: a background subtraction of raw the fluorescence image, a region of interest (ROI) segmentation, and a pixel-by-pixel ratio calculation within ROI using the intensities from 405 nm and 488 nm excitations (*I*_405_/*I*_488_). These processes were repeated at all timestamps to generate the final redox response trajectories. We estimated the background counts using an area without cells in each fluorescence micrograph under 405 nm and 488 nm excitation (*I*_405_ and *I*_488_). The background of raw experimental fluorescent images was subtracted to give the final background-corrected images for calculating the *I*_405_/*I*_488_ ratios. The region of interest (ROI) segmentation was achieved using the *k*-means segmentation approach to get the cellular boundary from the *I*_488_ image. After ROI segmentation, the ratiometric intensity for each extracellular treatment was obtained by calculating the mean value of ratiometric image (*I*_405_/*I*_488_) intensity pixel by pixel within ROI.

### Intracellular copper content quantification by ICP-MS Analysis

ICP-MS analysis was applied to quantify the copper content in cytosolic and mitochondrial fractions of HEK293 cells. We examined the averaged Cu contents in both compartments from three independent experiments. In short, the mitochondrial and cytosolic fractions were separated using the BioVision Mitochondria/Cytosol Fractionation Kit (Cat# K256-25, BioVision, Milpitas, CA, USA). The fractions were digested in 50 μL of ultrapure 70% nitric acid, heated at 95 °C for 2 hr in capped metal-free tubes, and gradually cooled down to room temperature to achieve complete mineralization. The samples were then further diluted in ultrapure water to a final 3% nitric acid concentration and analyzed by triple quadrupole inductively coupled plasma-mass spectrometry (Agilent 8800 ICP-QQQ). Copper concentrations were determined by using standard curves, and values were normalized to total sulfur content, as measured by ICP-MS on the same samples. We quantified the averaged Cu contents for cells that were under basal (i.e., no treatment), Cu-stressed (i.e., 200 μM CuCl_2_ for 1 hr), and Cu-depleted (i.e., 200 μM CuCl_2_ for 1 hr and followed by 200 μM BCS for another 1 hr) conditions.

### Proteomic measurement

Cytosolic and mitochondrial protein fractions were isolated from HEK293 cells under three treatment conditions: no treatment (Basal); 200 μM CuCl_2_ for 1 hr (Cu); and 200 μM CuCl_2_ for 1 hr followed by 200 μM BCS for another 1 hr (Cu→BCS) using the BioVision Mitochondria/Cytosol Fractionation Kit. The samples were added to an equal volume of 100 mM ammonium bicarbonate solution and heated at 95 ºC for 5 min. The reduction and alkylation reactions were carried out by adding dithiothreitol to a final concentration of 5 mM at 37 ºC for 1 hr, followed by adding iodoacetamide to a final concentration of 20 mM at room temperature for 30 min in the dark. Trypsin (1/40, w_trypsin_/w_protein_) was then added and incubated at 37 ºC overnight. The reaction was stopped by adding trifluoroacetic acid. The digested peptides were cleaned up using C18 Ziptips and vacuum dried using a CentriVap (Labconco). Each dried sample was resuspended in water with 0.1% FA for liquid chromatography-mass spectroscopy (LC-MS) analysis.

The nanoLC was coupled to a timsTOF Pro (Bruker Daltonics, Germany) via a CaptiveSpray source. Peptide samples were loaded onto an in-house packed column (75 μm x 25 cm, 1.9 μm ReproSil-Pur C18 particle (Dr. Maisch GmbH, Germany), column temperature 40 °C) with buffer A (0.1% FA in water) and buffer B (0.1% FA in ACN) as mobile phases. The total LC gradient is ∼21 min long. The concentration of buffer B increases from 2% to 30% during the first 17.8 min, further increases to 95% till 18.3 min, and maintains at 95% for another 2.4 min. A data-independent acquisition parallel accumulation-serial fragmentation (diaPASEF) scheme with 24 m/z and ion mobility windows was used. The electrospray voltage and the ion transfer tube temperature were 1.6 kV and 180 °C, respectively. Full MS scans were acquired over the m/z range of 150–1700. The collision energy was ramped linearly as a function of the mobility from 27 eV at 1/K_0_ = 0.85 Vscm^-2^ to 45 eV at 1/K_0_ = 1.3 Vscm^-2^.

The software Spectronaut v15 (Biosynosis, Switzerland) and an in-house spectral library were used for peptide and protein identification and quantification. Cysteine carbamidomethylation was listed as a fixed modification, and methionine oxidation and acetylation as variable modifications. The false discovery rate was <1% at both peptide and protein levels. Log2-transformation and median normalization were used to normalize the protein quantification data. Comparative analysis was performed using empirical Bayes moderated tests implemented in the R/Bioconductor limma package[21]. The limma statistical method was applied to statistically assess the difference in protein abundance between two experimental conditions from a smaller sample size per group (n = 3) [20]. Only proteins with an adjusted p-value below the threshold (alpha = 0.05) were considered statistically significant.

## Results

### Overview of experiments setup and imaging approach

To investigate the subcellular antioxidant defense, we imaged live HEK293 cells transfected with mito-Grx1-roGFP2 or cyto-Grx1-roGFP2 redox sensors using epifluorescence microscopy. Only cells with proper cellular morphology throughout the treatment and imaging steps were used to ensure physiological relevance. The scientific conclusion is drawn from a total of 107 cells (at least 3 cells for each condition).

To understand the subcellular redox responses, we treated HEK293 cells with oxidants (i.e., H_2_O_2_ or CuCl_2_) to induce oxidative stress, followed by reagents to remove the oxidants. (i.e., DTT or BCS). The redox status of mitochondria and cytosol under various oxidative and copper stress were examined by using a GSH/GSSG-based fluorescent protein sensor, Grx1-roGFP2. Grx1-roGFP2 specifically detects the glutathione redox couple’s ratio changes (i.e., GSH/GSSG ratio) through a fast equilibrium with the intracellular GSH/GSSG pool and serves as a fast intracellular redox sensor. The oxidized and reduced form of Grx1-roGFP2 exhibits maximum 520 nm fluorescence intensity under 405 nm and 488 nm excitation, respectively. A more oxidized environment will enrich the oxidized sensor and thus increase 405 nm and decrease 488 nm induced fluorescence intensity. Thus, the fluorescence intensity ratio between the two channels (i.e., *I*_405_/*I*_488_) can indicate the intracellular oxidation level.

The cells under various conditions (i.e., basal, CuCl_2_, and BCS) were further analyzed by liquid chromatography-mass spectroscopy. Proteomic results from mitochondria and cytosol under basal and Cu-stressed conditions were compared to identify proteins showing significant changes upon Cu treatments. Comparative analysis of proteins showing significant changes in two compartments reveals the mitochondria-specific redox-regulated proteins.

We transfected organelle-specific redox sensors, mito-Grx1-roGFP2 and cyto-Grx1-roGFP2, into HEK293 cells to probe the redox status at different organelles. Fig. 1A shows that the fluorescence images of the mito-Grx1-roGFP2 colocalize with a commercially available mitochondrial tracker, Mito ID. The fluorescence images of the cyto-Grx1-roGFP2 overlay nicely with the bright field micrograph of the cells (Fig. 1B). These results collectively demonstrate that mito-and cyto-Grx1-roGFP2 successfully targeted desired locations and confirm the feasibility of organelle-specific quantification of subcellular redox status.

**Figure 1.**
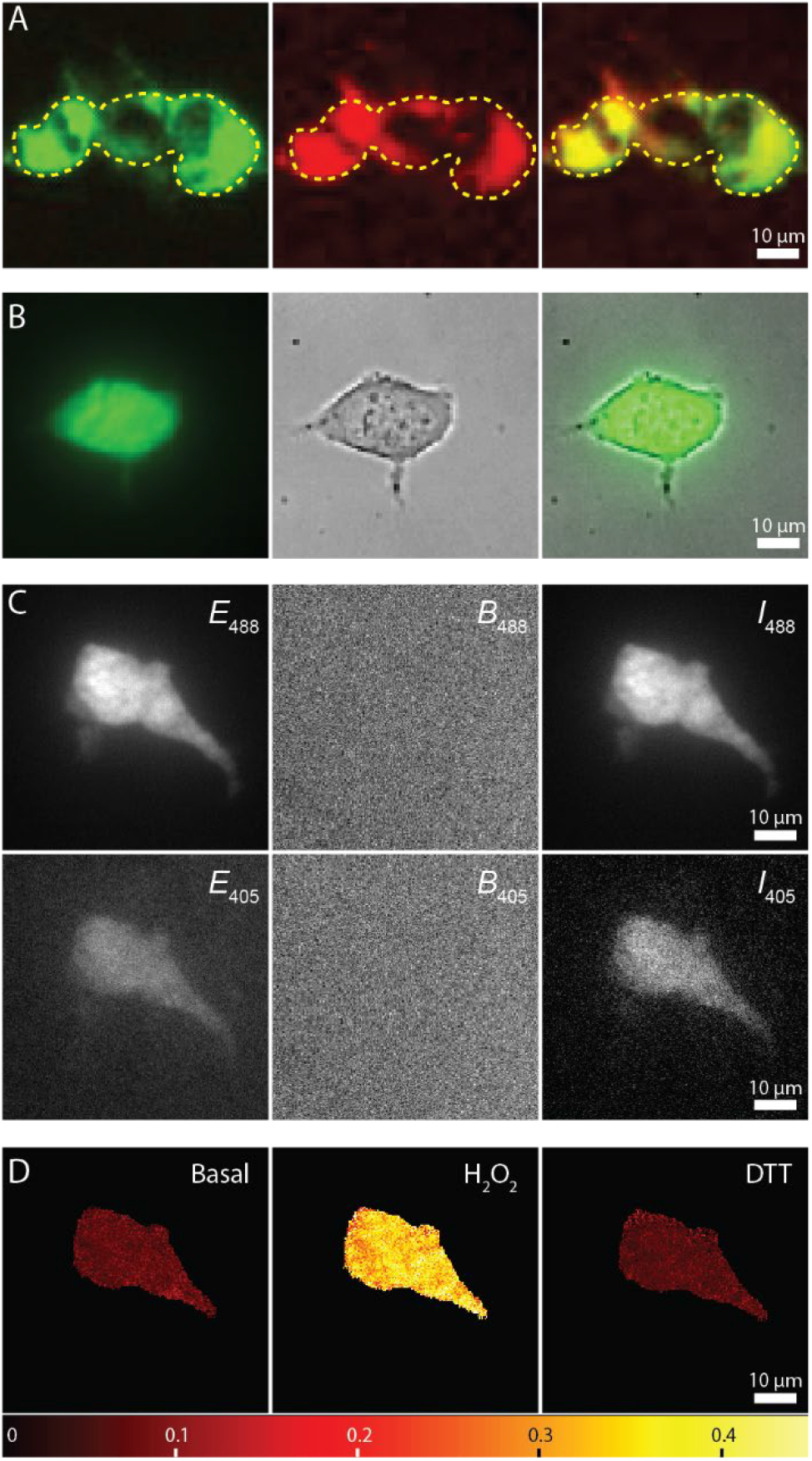
Organelle-specific redox sensing. (**A**) Fluorescence micrographs of a HEK293 cell show that mito-Grx1-roGFP2 (left) and mitochondria marker, MITO-ID, (center), are colocalized (right). (**B**) Similar as **A**, the cyto-Grx1-roGFP2 (left) and the cell transmission image (center) are well-aligned (right). (**C**) Image processing scheme to obtain background-removed fluorescence micrographs in 488 nm and 405 nm channels (left column: experimental images; center column: background images; right column: background-removed micrograph). The background-removed images were used for cell segmentation and ratiometic images. (**D**) ratiometric images under different conditions (basal, 200 μM H_2_O_2_, 1 mM DTT).

The redox status of subcellular compartments was estimated from the two-colored fluorescence images. Fig. 1C shows the exemplary imaging processing scheme to quantify cytosolic redox status under the basal condition. A cell transfected with the cyto-Grx1-roGFP2 sensor was imaged by epifluorescence microscopy under 405 nm and 488 nm excitations. The resulting raw images (*E*_405_ and *E*_488_) were corrected by the background image subtraction (*B*_405_ and *B*_488_) to generate the final fluorescence images (*I*_405_ and *I*_488_). The fluorescence intensity of each pixel within the ROI in the 405 nm channel was divided by the 488 nm channel. Averaging throughout the ROI gives the ratio (right column of Fig. 1C), reporting the subcellular redox status. The exemplary ratiometric images under basal, 200 μM H_2_O_2,_ and 1 mM DTT (Fig. 1D) show different ratios, indicating different cytosol redox statuses.

### Both cytosol and mitochondria show fast and robust responses to extracellular oxidative stresses

Several control experiments were first conducted to ensure that the oxidative responses are indeed induced by the oxidants (i.e., H_2_O_2_ and Cu) but not other reagents. As shown in Fig. 2A, we quantified the steady-state cellular redox states for HEK293 cells under basal (i.e., no treatment), 1 mM DTT, and 200 μM BCS treatments for extended imaging time (3 hours). Both mitochondria and cytosol under these three conditions show stable responses without significant changes throughout the imaging experiment. The results illustrate that these non-oxidants would not cause oxidative responses in our following experiments. We then examined the mitochondrial and cytosolic redox responses under sequential oxidative and reductive treatments to verify the fast responses of Grx1-roGFP2. HEK293 cells were treated with 200 μM H_2_O_2_ followed by 1 mM DTT and quantified the subcellular redox responses by the mito- and cyto-Grx1-roGFP2 sensors. Fluorescence micrographs were taken at different treatment times (varying from 2 to 90 min) to explore if the prolonged treatments have more profound effects on the cellular redox status. The left panel of Fig. 2B summarizes the redox readouts with 2 min treatment time. Both mitochondria and cytosol remain stable under basal conditions. Treatments with 200 μM H_2_O_2_ led to rapid (< 10 s) and significant increases in both mitochondria and cytosol. Switching from H_2_O_2_ to 1 mM DTT solution ameliorated oxidative stress in both compartments, with mitochondria showing a slightly slower response. Similar results were also observed when cells were challenged with a longer 90 min treatment time (right panel of Fig. 2B). These results are consistent with previous studies [7], indicating that the cellular machinery is highly effective to respond environmental oxidative stresses in both cytosol and mitochondria.

**Figure 2.**
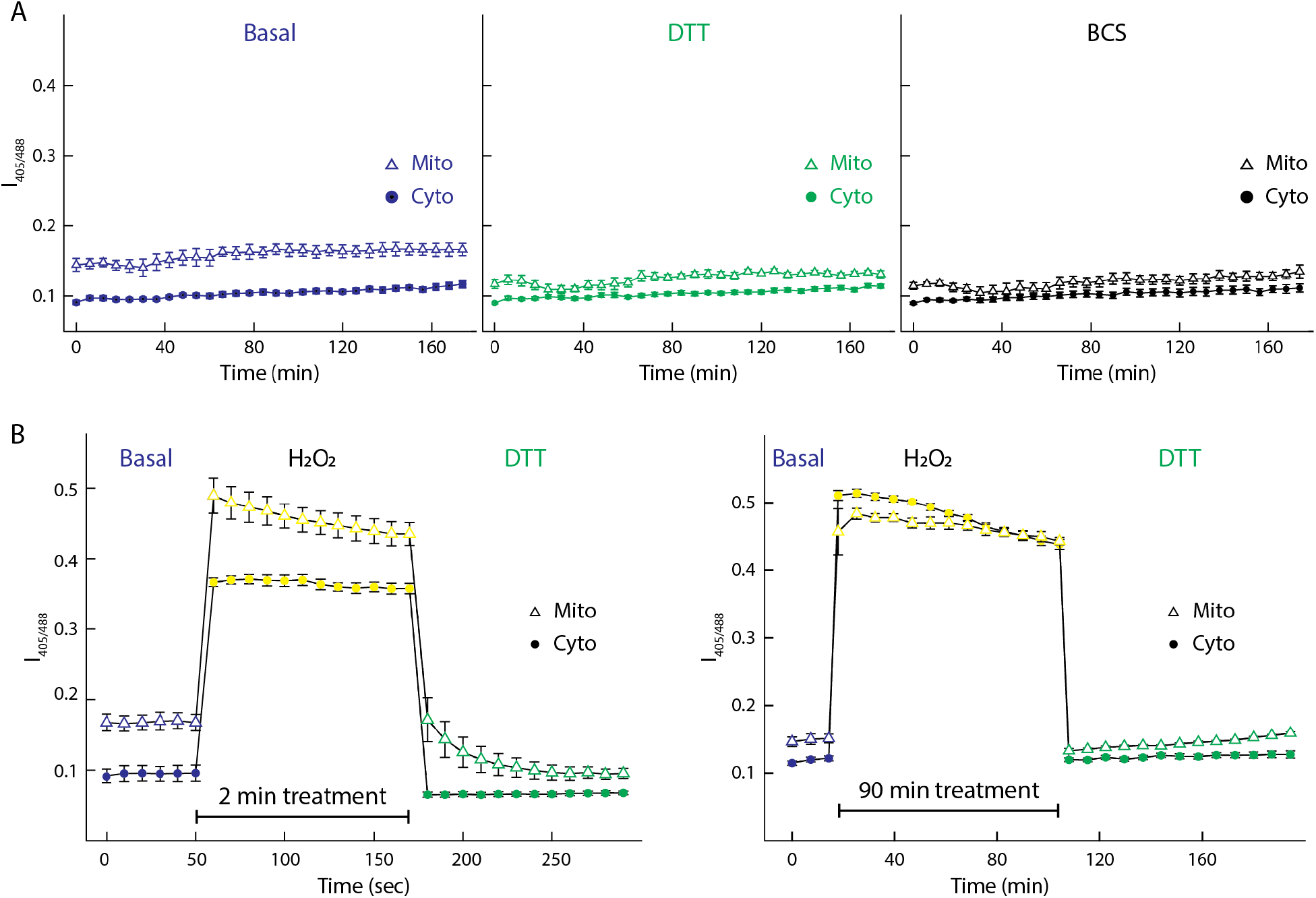
Subcellular redox responses under basal, DTT, BCS, and H_2_O_2_ treatments. (**A**) Control experiments of mitochondrial (triangles) and cytosolic (circles) redox responses. HEK 293 cells were under basal conditions (left) or treated with 1 mM DTT (center) or 200 μM BCS (right). No significant changes were observed in both compartments under these conditions. (**B**) Comparison of mitochondrial and cytosolic redox responses with 2 min (left) and 90 min (right) treatments. Both mitochondria and cytosol show rapid and robust redox responses with extracellular oxidative stress.

The fast and robust sensor system also provides a reliable platform to examine the Cu-induced oxidative stresses in cells.

### Mitochondria and cytosol handle Cu-induced oxidative stress differently

Using Grx1-roGFP2, we determine the mitochondrial and cytosolic oxidative status in real-time and investigate the effect of Cu on cellular antioxidative responses. Subcellular live-cell imaging experiments were conducted under sequential Cu-stressed (i.e., 200 μM CuCl_2_) and Cu-depleted (200 μM BCS) conditions with treatment times varying from 2 min to 90 min.

Fig. 3A summarizes the redox response results for cells with a short treatment time (< 15 min). Both mitochondrial and cytosolic oxidative states (i.e., *I*_405_/*I*_488_) remain stable throughout the experiment, indicating that transient Cu-relevant perturbations do not induce significant oxidative stress in both compartments under short Cu-treatment times. In contrast, consistent increase of oxidative stress was observed in both mitochondria and cytosol when the treatment time was longer than 35 min (Fig. 3B). The mitochondrial oxidative stress tends to increase faster than the cytosol and reaches a steady state after ∼40 min. When cells were treated with the cell-impermeable Cu chelating reagent, BCS, after Cu stimulation, mitochondria showed a quick response and thus a decrease of oxidative stress, and then gradually returned to the redox status similar to basal condition after ∼20 min. However, the oxidative stress in cytosol continuously increases during Cu treatment and unexpectedly keeps rising even after Cu wash-off by BCS (Fig. 3B and C).

**Figure 3.**
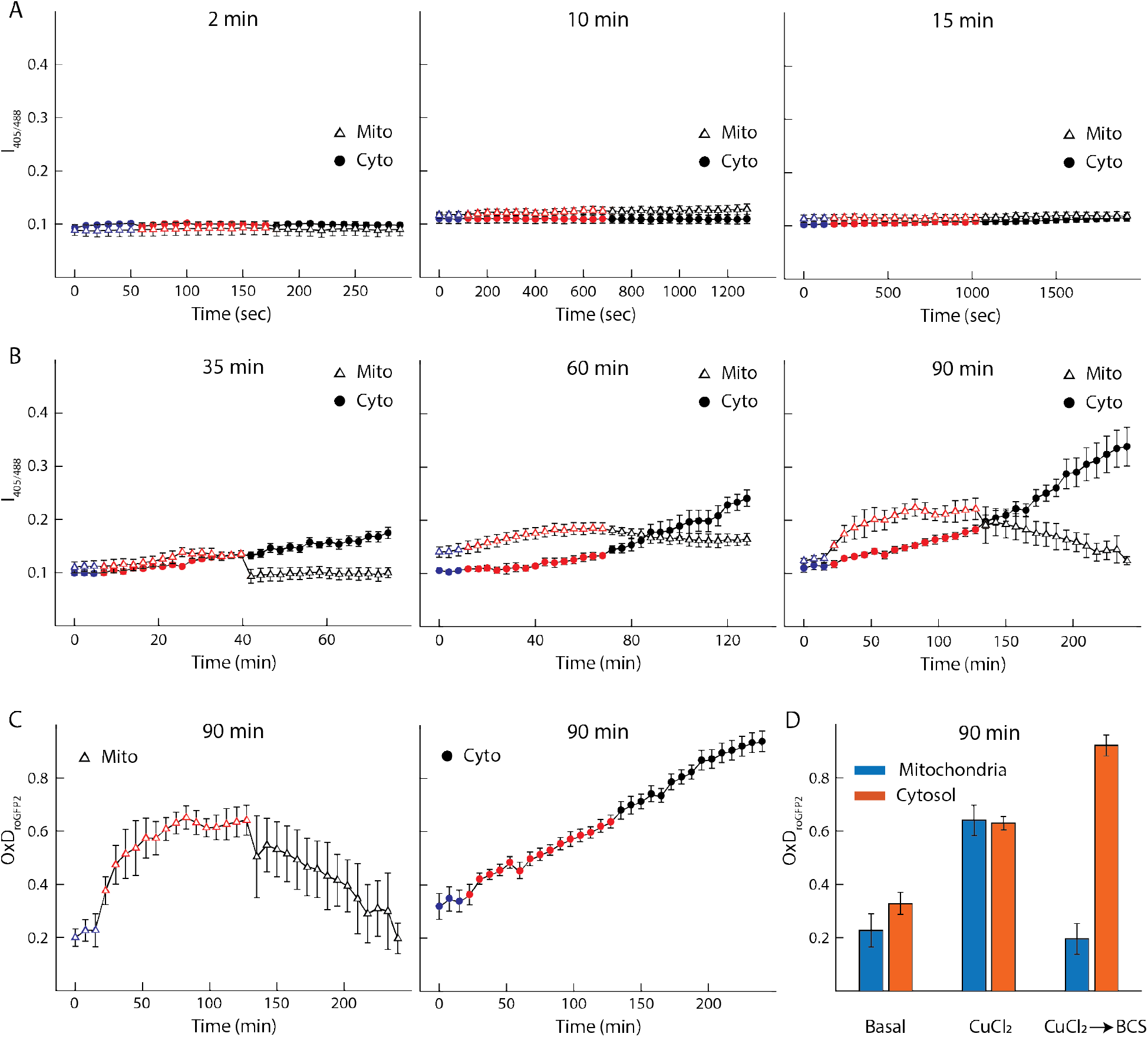
Subcellular Cu-induced redox responses under various Cu-stressed treatment times. (**A**) Mitochondrial and cytosolic redox status comparison of HEK293 cells transitioning from 2-15 min basal (blue curve), Cu-stressed (200 μM CuCl_2_, red curve), and Cu-deficient (200 μM BCS, black curve) conditions. (**B**) Same as **A**, but with longer treatment times (varying from 35-90 min). (**C**) Corresponding oxidation degree of mito and cyto-Grx1-roGFP2 (OxD_roGFP2_) and (**D**) summarized population bar graph. The OxD were calculated from the mean ratio at each time point from the data with 90 min treatment time.

To put our results into cellular context, we further calculated the corresponding oxidation degree of the redox sensor (OxD_roGFP2_) according to the equations described previously [20, 22]. Figure 3C shows the corresponding OxD_roGFP2_ results in mitochondria and cytosol under sequential 90 min CuCl_2_ and 90 min BCS treatments. The bar graph compares the value of OxD_roGFP2_ in mitochondria and cytosol at the end of each treatment stage (Fig. 3D). Mitochondria and cytosol have similar OxD_roGFP2_ under basal (i.e., mito: 23% vs. cyto: 33%) and Cu-stressed (i.e., mito: 64% vs. cyto: 63%) conditions. But upon BCS treatment, cytosolic OxD_roGFP2_ further increased to 92%, while mitochondrial OxD_roGFP2_ returned back to a basal-like level.

To further understand this intriguing observation, we performed subcellular live-cell imaging with three-hour 200 μM CuCl_2_ long treatments. As expected, the mitochondrial GSSG populations increase much faster than the cytosol but eventually reach a plateau after ∼ 40 min (Fig. 4A). The plateau value (∼0.25) is much lower than that (∼0.5) of the H_2_O_2_ treatments, suggesting that mitochondria effectively maintain the ROS to a level that does not require fully bleach GSH. The cytosolic GSSG level shows a slow but steady increase (Fig. 4A). Interestingly, when we treated the cells with 35 min of CuCl_2_ and switched media to BCS solution for an extended period of time, the cytosolic GSSG levels still showed a similar growth trend as in the pure CuCl_2_ solution (Fig. 4B).

**Figure 4.**
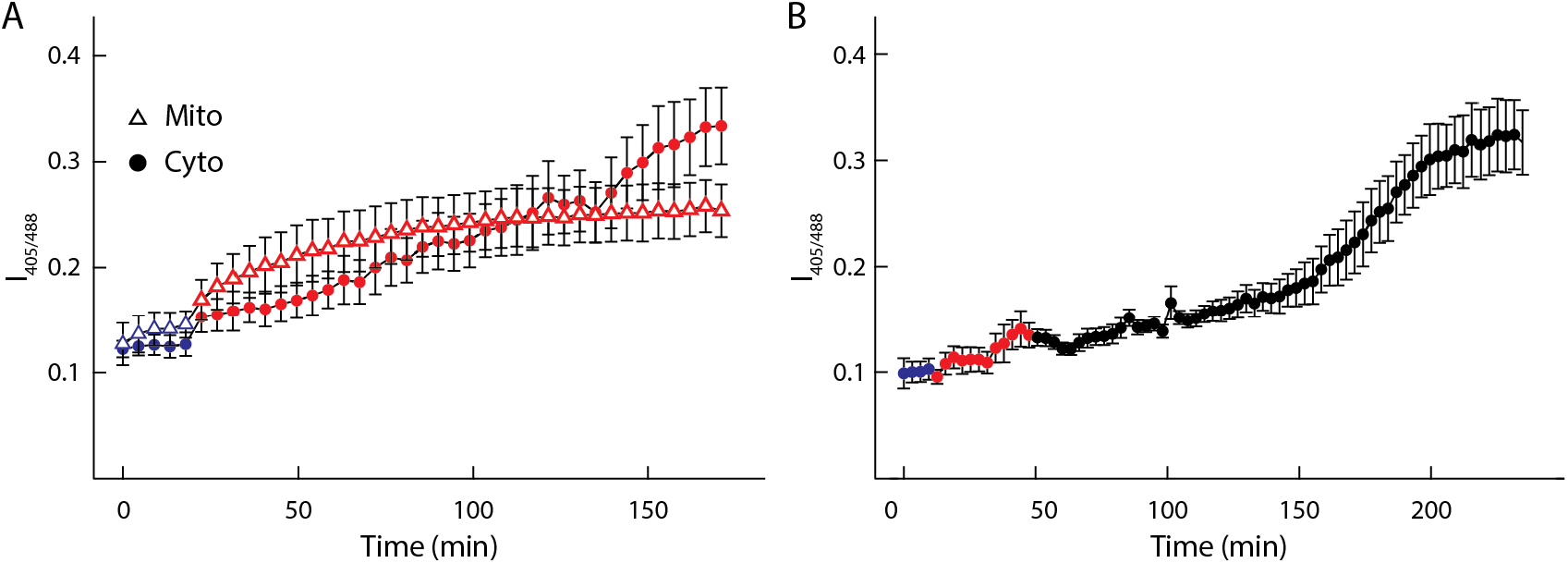
Subcellular redox responses under prolonged treatments. (**A**) Mitochondrial and cytosolic redox status under 3 hr Cu treatments. (**B**) Similar to **A** but with 35 min Cu treatment followed by 3 hr BCS treatments.

Our real-time redox quantification results clearly show that Cu-induced oxidative stress happens at much slower rates than H_2_O_2,_ and the prolonged Cu treatment induces significant GSSG/GSH changes. Besides, the differential redox responses between mitochondria and cytosol suggest that mitochondria have a more rapid reaction mechanism than cytosol. These observations further raise the following questions: Are the fast mitochondrial Cu-responses due to Cu preferentially imported into the mitochondria? Does the fast mitochondrial BCS responses indicate that the mitochondria have other antioxidant mechanisms than the cytosol? If so, who will be the key candidates? To answer these questions, we use two mass spectrometry approaches to quantify both the subcellular Cu content and potential proteins responsible for the more rapid antioxidative defense in mitochondria.

### Mitochondria modulate Cu levels more efficiently than the cytosol

To quantify the copper content in both mitochondria and cytosol under different Cu-related oxidative conditions, we performed the subcellular ICP-MS analysis. Fig. 5A shows the averaged Cu content (i.e., Cu/S) in mitochondria and cytosol obtained from cells under basal (i.e., no treatment), Cu-stressed (i.e., 200 μM CuCl_2_ for 1 hr), and Cu-depleted (i.e., 200 μM CuCl_2_ for 1 hr and followed by 200 μM BCS for another 1 hr) conditions. The amount of Cu was normalized by the amount of S for comparisons since intracellular S is equally distributed in the cell [23, 24]. Under Cu-stressed conditions, mitochondria underwent a significant Cu content elevation with a final Cu content ∼ 6 fold of the cytosol. Upon switching to BCS, the Cu content decreased in both compartments. Despite having a higher Cu content after Cu treatment, the mitochondria also demonstrate a more efficient Cu efflux after BCS treatment. The averaged Cu efflux mitochondrial effluxed Cu concentration (i.e., ΔCu/S_mito_) is ∼ 7 fold higher than the cytosol (i.e., ΔCu/S_cyto_, Fig. 5A).

**Figure 5.**
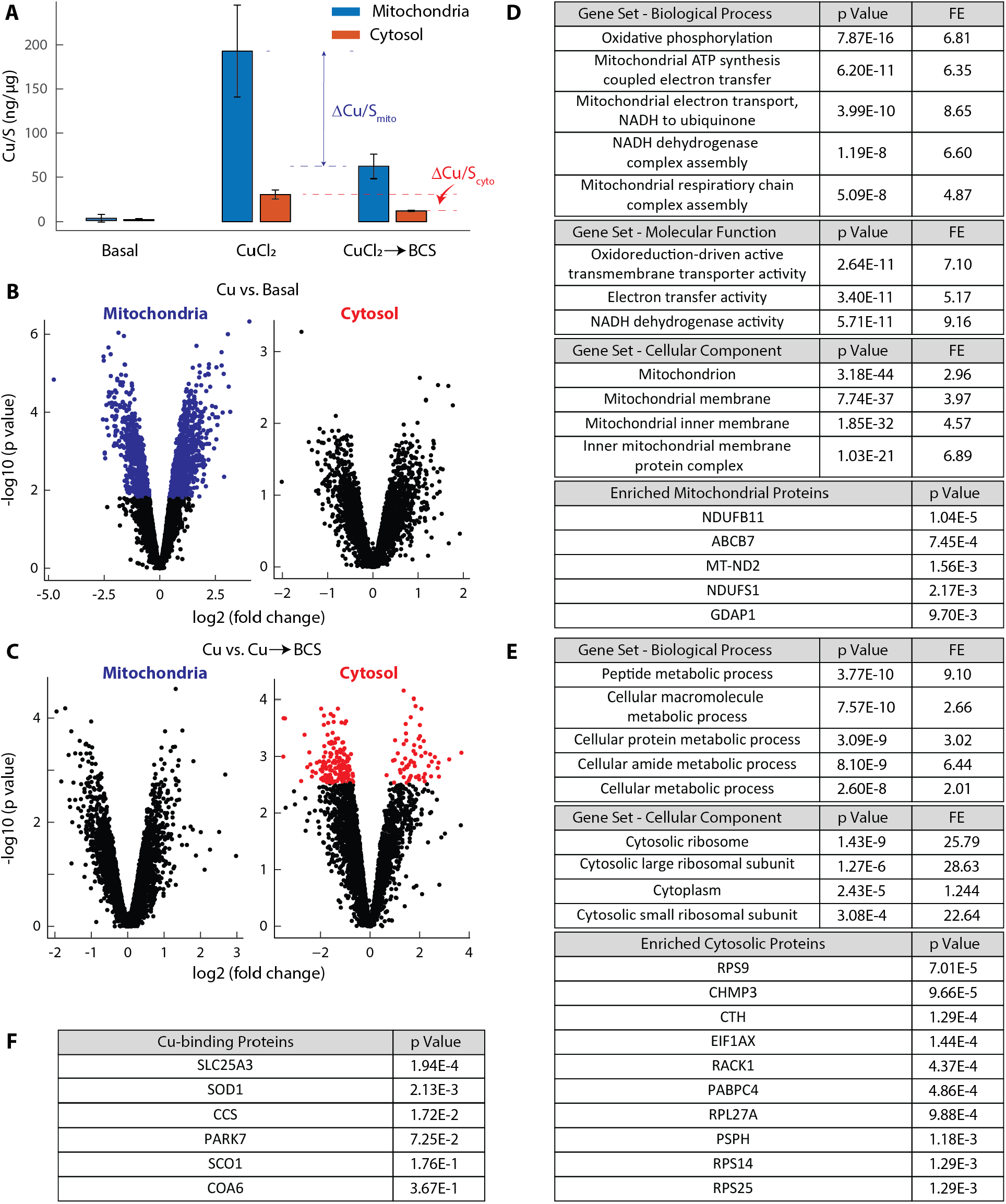
Subcellular ICP-MS and GO analysis results. (**A**) Relative mitochondrial and cytosolic Cu concentrations from ICP-MS measurements under different treatments. (**B**) Mitochondrial (left) and cytosolic (right) volcano plots when cells transition from basal to Cu-stressed conditions. (**C**) Same as **B** but for cells transition from Cu-stressed to Cu-deficient conditions. (**D**) Mitochondrial GO analysis results reveal the key responding biological process, cellular compartment, and proteins. Here, fold enrichment (FE) is defined as the ratio of sample frequency representing the number of genes inputted that are involved in a GO enriched term and the background frequency of total genes annotated to that same term. (**E**) Same as **D** but for the cytosol. (**F**) GO analysis results for proteins responsible for cellular Cu homeostasis.

### Mitochondria and cytosol respond differently to Cu-induced oxidative stress at the protein level

Proteomic profiling of mitochondria and cytosol was carried out to identify potential redox-regulated proteins that lead to differential Cu responses between these two compartments. The proteins with significant abundance changes in the mitochondria and cytosol upon CuCl_2_ and BCS treatments are statistically estimated and visualized by volcano plots (Fig. 5B and C). Under Cu stress, several proteins in the mitochondria, but not cytosol, underwent significant abundance changes (Fig. 5B). However, the subsequent removal of Cu only resulted in substantial protein abundance changes in the cytosol (Fig. 5C).

Gene ontology (GO) enrichment analysis [25] identified redox-driven active transmembrane activity and electron transfer activity as the top upregulated molecular functions (Fig. 5D). Furthermore, upregulated proteins with significant abundance changes identified in these two top-scoring molecular functions are the subunit of either the complex IV cytochrome *c* oxidase (COX5A, COX5B, COX7A2, COX6B1, MT-CO2) or complex I NADH ubiquinone/ubiquinone oxidoreductase complex (MT-ND2, MT-ND3, NDUFAF2, NDUFA10, NDUFA9). These enzyme complexes are known to be responsible for mitochondrial oxidative phosphorylation (OXPHOS). It is worth noting that the identified proteins in these two complexes are not Cu-binding proteins, indicating the enriched pathways likely do not involve handling Cu directly but digesting the effects induced by the Cu. Our observation of increased complex IV proteins with higher Cu content is consistent with a recent report [26]. In addition, our GO analysis further provides insights into the key subunits contributing to the observations.

At the protein level, we also use the GO resource to identify the enriched mitochondrial proteins under Cu treatments. Fig. 5D lists the top 5 of the 159 mitochondrial proteins that show significant abundance changes and are relevant to the redox-regulated functions. The NDUFB11, MT-ND2, and NDUFS1 are key proteins involved in the NADH: ubiquinone reductase complex assembly [27]. ABCB7 is involved in transporting heme from the mitochondria to the cytosol and potentially plays a role in iron homeostasis. Interestingly, GDAP1, a protein that belongs to the GSH S-transferases (GSTs) enzyme family, also presents as one of the top 5 proteins. GDAP1 displays GSH-dependent activity. Studies suggest a protective role of GDAP1 against oxidative stress related to the intracellular levels in GSH [28, 29]. GSH is downregulated in fibroblasts from patients with mutations in GDAP1 alleles. Therefore, it is suggested that GDAP1 protects mitochondria by reducing the ROS by modulating the GSH/GSSG equilibrium.

A similar analysis was also performed to identify cellular pathways that were enriched in the cytosol under BCS treatments (Fig. 5E). The GO analysis identified the peptide metabolic process as the most significantly enriched and upregulated biological process and the ribosome as the main cellular compartment. In fact, the enriched cytosolic proteins, RPS9, RPL27A, RPS14, and RPS25 belong to the cytosolic ribosomal protein family. EIF1AX and PABPC4 are also involved in the translation process. Collectively, these data indicate that the HEK293 cells, after 1 hr BCS treatments, were still actively synthesizing proteins or peptides to respond to the Cu-related changes. It’s noteworthy that the glutathione metabolic process belongs to the peptide metabolic process, which is highly relevant to our experiment due to the role of glutathione as the antioxidant during this Cu-induced oxidative stress response. Interestingly, CTH and PSPH are another two significantly upregulated proteins. CTH is one of the two major pathways to generate cysteine, which is the major resource to generate GSH in the cytosol. PSPH, on the other hand, catalyzes the last irreversible step in the biosynthesis of L-serine, which can be used in glutathione synthesis.

It is well-accepted that a change in cellular redox status can modulate the redox state of metalloproteins and thus lead to Cu redistribution to different cellular compartments. It is also known that excess intracellular Cu can perturb cellular redox balance [30, 31]. Here we compared the subcellular redox responses with the GO data to understand the effects of Cu on subcellular redox responses. Using the proteomic data, we first searched mitochondrial Cu-binding proteins [32] that show significant abundance changes. Among the major Cu-binding mitochondrial proteins, including SLC25A3, CCS, SOD1, SCO1, COA6, and PARK7, only SLC25A3 and SOD1 were identified with significant abundance changes upon 1hr Cu treatment in mitochondria (Fig. 5F). SLC25A3 is a protein that transports the Cu-ligand complex across the inner membrane for Cu storage in the mitochondrial matrix. The upregulated SLC25A3 potentially channels the excess Cu to the matrix and minimizes the Cu toxicity [33]. In contrast, the abundance of SOD1 was significantly decreased upon Cu treatment but gradually increased back after the BCS addition.

## Discussion

Sensing and understanding subcellular Cu-dependent antioxidative defense is important for preventing Cu-induced redox damage due to reactive oxygen species. We reported here that mitochondria and cytosol show different antioxidative responses upon Cu-dependent oxidative stress. In this work, by using combined approaches of organelle-specific redox sensor Grx1-roGFP2 and non-targeted proteomics, we have studied the mitochondrial and cytosolic Cu-induced redox response with different treatment durations. Compared to the H_2_O_2_-induced responses, subcellular Cu-induced oxidative stress happens with much slower rates and fewer extents. Furthermore, GO analysis suggests that mitochondria react more promptly to Cu-dependent treatments than the cytosol and boost complex I, complex IV, and GSH relevant proteins expressions; cytosol, on the other hand, mainly increases GSH-related proteins expression.

The tripeptide glutathione (reduced form: GSH; oxidized disulfide form: GSSG) is one of the major players in maintaining intracellular redox equilibrium by altering the GSH/GSSG ratio [34]. Redox-sensitive Grx1-roGFP2 is a quantitative, ratiometric, and pH-independent [35] redox sensor that visualizes cellular redox status changes in real time [2]. Grx1 enforces continuous fast equilibration between the two redox couples (i.e., roGFP2_red_/roGFP2_ox_ and 2GSH/GSSG) through the monothiol mechanism [36, 37], enabling the detection of small oxidative stress changes of high concentration of GSH. Under oxidative stress, the disulfide bond formed on roGFP2 promotes the protonation of the GFP chromophore, shifting the excitation peak from 488 nm to near 405 nm and enabling the quantification of oxidation levels using the fluorescence ratio under 405 nm and 488 nm excitations.

In other words, the observed Cu-induced differential redox responses between mitochondria and cytosol report the GSH/GSSG ratio changes, which likely originated from the Cu-induced ROS. This ROS generation step (Fig. 3B, 90 min Cu data) is relatively slow compared to the H_2_O_2_ induced oxidative stress (Fig. 2B, 90 min H_2_O_2_ data). With treatment times longer than 15 min, mitochondria show a more rapid increase of GSSG populations upon Cu stress and a decrease upon BCS treatment (Fig. 3B and 3C). In contrast, cytosol shows a gradually increased GSSG population after Cu treatment, and the population continues to grow even with BCS treatment (Fig. 4B). Since prolonged BCS treatments do not alter the GSH/GSSG ratio (Fig. 2A), the observed increasing oxidative stress must be originated from the prolonged Cu treatments. This observation implies that even though the GSSG population growth is originated from a Cu-initiated process, subsequent Cu removal can’t slow down the process that eventually leads to the GSSG generation.

It is well accepted that mitochondria are one of the most important compartments mediating Cu-induced hepatotoxicity [38]. The faster mitochondrial Cu response suggests that Cu preferentially targets the mitochondria once entering the cellular environment through an unknown pathway. Although the organelle-specific ICP-MS results (Fig. 5A) can’t provide kinetic information on Cu uptake, the higher Cu content in the mitochondrial also supports the same concept. Previous studies have demonstrated that Cu treatments induce a rise in mitochondrial ROS formation, membrane potential decline, and cytochrome *c* expulsion. These phenotypes result from Cu’s disruptive effect on the mitochondrial respiratory chain [39].

The GO analysis of Cu-stressed mitochondria highlights that OXPHOS complex I (i.e., NADH: ubiquinone reductase) and IV (i.e., cytochrome *c* oxidase) are the two most upregulated populations. It has been shown that cytochrome c oxidase activities collapse with increasing Cu concentration, and mitochondrial H_2_O_2_ production is significantly elevated when mitochondria are treated with a complex I inhibitor. The upregulated complex I and IV protein levels are likely a compensation mechanism to maintain OXPHOS functions. Further, GSH is produced exclusively in the cytosol and actively pumped into mitochondria [40]. Cu has been shown to decrease mitochondrial GSH levels [39]. In contrast, overexpression of the mitochondria outer membrane protein Gdap1 has been shown to increase the total cellular glutathione level and protect against endogenous oxidative stress caused by glutathione depletion [28]. Our GO analysis reveals that GDAP1 indeed is upregulated upon Cu stressed conditions. Since the GSSG formed in mitochondria is unlikely to exit this organelle [41], upregulated GDAP1 would potentially further bring more GSH into mitochondria and minimize the ROS damage. In fact, the fast mitochondrial BCS response suggests that mitochondria restore the basal GSH/GSSG ratio more efficiently than the cytosol once the Cu is reduced, indicating mitochondria are most likely to utilize multiple means to minimize ROS damages. Previous studies reported that mitochondria have more tightly controlled total and labile Cu pools than the whole cell, therefore are better at regulating mitochondria Cu [42]. Our results suggest that mitochondria also have multiple ROS regulatory pathways (i.e., GSH increments and upregulated OXPHOS functions). The additive effects of the superior antioxidant defenses and Cu controls ultimately lead to the observed mitochondrial behaviors.

It is interesting to observe mitochondrial Cu-binding proteins SOD1 and SLC25A3 changing their abundance. The delivery of Cu to mitochondria is obligatory for cell survival due to the essential role of Cu for cytochrome c oxidase activity [32, 43-46]. SOD1, localized to the intermembrane space (IMS), is the front-line antioxidant enzyme that catalyzes the disproportionation of superoxide radicals and integrates oxygen availability to redox regulate NADPH production [47]. We reason the observed decreased mitochondrial SOD1 abundance to the lack of apo-SOD1 polypeptide source under Cu-stressed conditions since only the very immature form of the SOD1 polypeptide that is apo for both Cu and Zn can efficiently enter mitochondria [48]. Under Cu stress, the population of the SOD1 apo polypeptide is likely to decrease significantly. SLC25A3 is a mitochondrial copper transporter required for cytochrome c oxidase biogenesis [33]. It has been shown that Slc25a3^—/—^ cells had reduced levels of the cytosolic copper enzyme SOD1 and its chaperone CCS [33]. Therefore, the upregulated SLC25A3 protein abundance can’t be responsible for the decreased mitochondrial SOD1 level. The upregulated SLC25A3 most likely transports the excess Cu to the matrix and minimizes the Cu toxicity [33].

## Conclusions

The interplay between Cu homeostasis and antioxidant defense is essential for many biological processes. Misregulation of one perturbs the other and is often observed simultaneously in neurodegenerative diseases. This study utilizes organelle-specific redox sensors to follow the real-time evolutions of Cu-induced oxidative stress in mitochondria and cytosol under basal, Cu-stressed, and Cu-deficient conditions. Non-targeted proteomics was also applied to identify proteins that undergo significant abundance changes. We found that mitochondria are more sensitive to Cu treatment than the cytosol and initiate the upregulations of the GSH productions and OXPHOS functions. In contrast, the cytosol responds to the Cu stress much slower. The Cu-induced oxidative stress continues to increase and reach a new steady state even after the extracellular Cu removal. Although this study focuses on the redox and protein abundance quantification in mitochondria and cytosol under various Cu stresses, a similar approach can be applied to examine the Cu-induced ROS in other organelles such as the endoplasmic reticulum and Golgi apparatus. Furthermore, the recent advance in super-resolution microscopy enables researchers to quantify protein kinetics and oligomeric states from a single-molecule perspective [49-53]. Considering Cu trafficking is mediated by the specific protein interaction, examining the protein interactions between Cu-binding proteins will further elucidate the Cu homeostasis mechanisms at different organelles. Overall, our findings provide insights into how redox and Cu homeostasis interplay by modulating specific protein expressions at the subcellular levels, shedding light on understanding the effects of Cu-induced redox misregulation on the diseases.

## Reference

1. Bertini I. Biological inorganic chemistry: structure and reactivity. University Science Books, 2007

2. Turski ML, Thiele DJ. New roles for copper metabolism in cell proliferation, signaling, and disease. Journal of Biological Chemistry 2009;284(2):717–21

3. Hatori Y, Yan Y, Schmidt K et al. Neuronal differentiation is associated with a redox-regulated increase of copper flow to the secretory pathway. Nature communications 2016;7:10640

4. Ogra Y, Tejima A, Hatakeyama N et al. Changes in intracellular copper concentration and copper-regulating gene expression after PC12 differentiation into neurons. Sci Rep 2016;6:33007

5. Kaler SG. Inborn errors of copper metabolism. Handb Clin Neurol 2013;113:1745

6. Graper ML, Huster D, Kaler SG et al. Introduction to human disorders of copper metabolism. Annals of the New York Academy of Sciences 2014;1314(1)

7. Gutscher M, Pauleau A-L, Marty L et al. Real-time imaging of the intracellular glutathione redox potential. Nature methods 2008;5(6):553–9

8. Schafer FQ, Buettner GR. Redox environment of the cell as viewed through the redox state of the glutathione disulfide/glutathione couple. Free radical biology and medicine 2001;30(11):1191–212

9. Meredith MJ, Reed D. Status of the mitochondrial pool of glutathione in the isolated hepatocyte. Journal of Biological Chemistry 1982;257(7):3747–53

10. Lu SC. Regulation of glutathione synthesis. In: Stadtman ER, Chock PBs (eds). Current Topics in Cellular Regulation: Academic Press, 2001, 95–116

11. Lutsenko S. Dynamic and cell-specific transport networks for intracellular copper ions. Journal of cell science 2021;134(21):jcs240523

12. Cannon MB, Remington SJ. Re-engineering redox-sensitive green fluorescent protein for improved response rate. Protein science 2006;15(1):45–57

13. Schymanski EL, Singer HP, Slobodnik J et al. Non-target screening with high-resolution mass spectrometry: critical review using a collaborative trial on water analysis. Analytical and bioanalytical chemistry 2015;407(21):6237–55

14. Mi H, Muruganujan A, Huang X et al. Protocol Update for large-scale genome and gene function analysis with the PANTHER classification system (v. 14.0). Nature protocols 2019;14(3):703–21

15. Garza NM, Griffin AT, Zulkifli M et al. A genome-wide copper-sensitized screen identifies novel regulators of mitochondrial cytochrome c oxidase activity. J Biol Chem 2021;296:100485. doi: 10.1016/j.jbc.2021.100485

16. Gao Q, Zhu H, Dong L et al. Integrated proteogenomic characterization of HBV-related hepatocellular carcinoma. Cell 2019;179(2):561-77. e22

17. Mills RJ, Titmarsh DM, Koenig X et al. Functional screening in human cardiac organoids reveals a metabolic mechanism for cardiomyocyte cell cycle arrest. Proceedings of the National Academy of Sciences 2017;114(40):E8372–E81

18. Stukalov A, Girault V, Grass V et al. Multilevel proteomics reveals host perturbations by SARS-CoV-2 and SARS-CoV. Nature 2021;594(7862):246–52

19. Patel NM, Moore JD, Blackwell HE et al. Identification of unanticipated and novel N-acyl L-homoserine lactones (AHLs) using a sensitive non-targeted LC-MS/MS method. PloS one 2016;11(10):e0163469

20. Morgan B, Sobotta MC, Dick TP. Measuring EGSH and H2O2 with roGFP2-based redox probes. Free Radical Biology and Medicine 2011;51(11):1943–51

21. Ritchie ME, Phipson B, Wu D et al. limma powers differential expression analyses for RNA-sequencing and microarray studies. Nucleic acids research 2015;43(7):e47–e

22. Meyer AJ, Dick TP. Fluorescent protein-based redox probes. Antioxidants & redox signaling 2010;13(5):621–50

23. Gräfenstein A, Rumancev C, Pollak R et al. Spatial Distribution of Intracellular Ion Concentrations in Aggregate-Forming HeLa Cells Analyzed by μ-XRF Imaging. ChemistryOpen 2022;11(4):e202200024. doi: 10.1002/open.202200024

24. Finney L, Mandava S, Ursos L et al. X-ray fluorescence microscopy reveals large-scale relocalization and extracellular translocation of cellular copper during angiogenesis. Proceedings of the National Academy of Sciences 2007;104(7):2247–52. doi: doi:10.1073/pnas.0607238104

25. Consortium GO. The gene ontology resource: 20 years and still GOing strong. Nucleic acids research 2019;47(D1):D330–D8

26. Ruiz LM, Jensen EL, Rossel Y et al. Non-cytotoxic copper overload boosts mitochondrial energy metabolism to modulate cell proliferation and differentiation in the human erythroleukemic cell line K562. Mitochondrion 2016;29:18–30

27. Signes A, Fernandez-Vizarra E. Assembly of mammalian oxidative phosphorylation complexes I–V and supercomplexes. Essays in biochemistry 2018;62(3):255–70

28. Noack R, Frede S, Albrecht P et al. Charcot–Marie–Tooth disease CMT4A: GDAP1 increases cellular glutathione and the mitochondrial membrane potential. Human molecular genetics 2012;21(1):150–62

29. López Del Amo V, Seco-Cervera M, García-Giménez JL et al. Mitochondrial defects and neuromuscular degeneration caused by altered expression of Drosophila Gdap1: implications for the Charcot–Marie–Tooth neuropathy. Human molecular genetics 2015;24(1):21–36

30. Hatori Y, Lutsenko S. The role of copper chaperone Atox1 in coupling redox homeostasis to intracellular copper distribution. Antioxidants 2016;5(3):25–40

31. Hatori Y, Lutsenko S. An expanding range of functions for the copper chaperone/antioxidant protein Atox1. Antioxidants & redox signaling 2013;19(9):945–57

32. Ruiz LM, Libedinsky A, Elorza AA. Role of Copper on Mitochondrial Function and Metabolism. Front Mol Biosci 2021;8:711227. doi: 10.3389/fmolb.2021.711227

33. Boulet A, Vest KE, Maynard MK et al. The mammalian phosphate carrier SLC25A3 is a mitochondrial copper transporter required for cytochrome c oxidase biogenesis. Journal of Biological Chemistry 2018;293(6):1887–96

34. Dooley CT, Dore TM, Hanson GT et al. Imaging dynamic redox changes in mammalian cells with green fluorescent protein indicators. Journal of Biological Chemistry 2004;279(21):22284–93

35. Schwarzländer M, Fricker M, Müller C et al. Confocal imaging of glutathione redox potential in living plant cells. Journal of microscopy 2008;231(2):299–316

36. Peltoniemi MJ, Karala A-R, Jurvansuu JK et al. Insights into deglutathionylation reactions: Different intermediates in the glutaredoxin and protein disulfide isomerase catalyzed reactions are defined by the γ-linkage present in glutathione. Journal of Biological Chemistry 2006;281(44):33107–14

37. Fernandes AP, Holmgren A. Glutaredoxins: glutathione-dependent redox enzymes with functions far beyond a simple thioredoxin backup system. Antioxidants and Redox Signaling 2004;6(1):63–74

38. Mehta R, Templeton DM, O’Brien PJ. Mitochondrial involvement in genetically determined transition metal toxicity: II. Copper toxicity. Chemico-biological interactions 2006;163(1-2):77–85

39. Hosseini M-J, Shaki F, Ghazi-Khansari M et al. Toxicity of copper on isolated liver mitochondria: impairment at complexes I, II, and IV leads to increased ROS production. Cell biochemistry and biophysics 2014;70(1):367–81

40. Lu SC. Glutathione synthesis. Biochim Biophys Acta 2013;1830(5):3143–53. doi: 10.1016/j.bbagen.2012.09.008

41. Olafsdottir K, Reed DJ. Retention of oxidized glutathione by isolated rat liver mitochondria during hydroperoxide treatment. Biochimica et Biophysica Acta (BBA)-General Subjects 1988;964(3):377–82

42. Dodani SC, Leary SC, Cobine PA et al. A targetable fluorescent sensor reveals that copper-deficient SCO1 and SCO2 patient cells prioritize mitochondrial copper homeostasis. Journal of the American Chemical Society 2011;133(22):8606–16

43. Cobine PA, Moore SA, Leary SC. Getting out what you put in: Copper in mitochondria and its impacts on human disease. Biochimica Et Biophysica Acta (BBA)-Molecular Cell Research 2021;1868(1):118867

44. Wen M-H, Xie X, Huang P-S et al. Crossroads between membrane trafficking machinery and copper homeostasis in the nerve system. Open Biology 2021;11(12):210128. doi: doi:10.1098/rsob.210128

45. Baker ZN, Cobine PA, Leary SC. The mitochondrion: a central architect of copper homeostasis. Metallomics 2017;9(11):1501–12. doi: 10.1039/c7mt00221a

46. Ohrvik H, Aaseth J, Horn N. Orchestration of dynamic copper navigation - new and missing pieces. Metallomics 2017;9(9):1204–29. doi: 10.1039/C7MT00010C

47. Montllor-Albalate C, Kim H, Thompson AE et al. Sod1 integrates oxygen availability to redox regulate NADPH production and the thiol redoxome. Proceedings of the National Academy of Sciences 2022;119(1):e2023328119. doi: doi:10.1073/pnas.2023328119

48. Field LS, Furukawa Y, O’Halloran TV et al. Factors controlling the uptake of yeast copper/zinc superoxide dismutase into mitochondria. Journal of Biological Chemistry 2003;278(30):28052–9

49. Pan M, Zhang Y, Yan G et al. Dissection of interaction kinetics through single-molecule interaction simulation. Analytical Chemistry 2020;92(17):11582–9

50. Chen H, Xie X, Chen T-Y. Single-molecule microscopy for in-cell quantification of protein oligomeric stoichiometry. Current Opinion in Structural Biology 2020;66:112–8

51. Santiago AG, Chen T-Y, Genova LA et al. Adaptor protein mediates dynamic pump assembly for bacterial metal efflux. Proceedings of the National Academy of Sciences 2017;114(26):6694–9. doi: 10.1073/pnas.1704729114

52. Chen T-Y, Santiago AG, Jung W et al. Concentration-and chromosome-organization-dependent regulator unbinding from DNA for transcription regulation in living cells. Nature communications 2015;6:7445

53. Chen T-Y, Jung W, Santiago AG et al. Quantifying multistate cytoplasmic molecular diffusion in bacterial cells via inverse transform of confined displacement distribution. The Journal of Physical Chemistry B 2015;119(45):14451–9

